# On the (Im)possibility to Reconstruct Plasmids from Whole Genome Short-Read Sequencing Data

**DOI:** 10.1101/086744

**Authors:** Sergio Arredondo-Alonso, Willem van Schaik, Rob J. Willems, Anita C. Schürch

## Abstract

Plasmids are autonomous extra-chromosomal elements in bacterial cells that can carry genes that are important for bacterial survival. To benchmark algorithms for automated plasmid sequence reconstruction from short read sequencing data, we selected 42 publicly available complete bacterial genome sequences which were assembled by a combination of long- and short-read data. The selected bacterial genome sequence projects span 12 genera, containing 148 plasmids. We predicted plasmids from short-read data with four different programs (PlasmidSPAdes, Recycler, cBar and PlasmidFinder) and compared the outcome to the reference sequences.

PlasmidSPAdes reconstructs plasmids based on coverage differences in the assembly graph. It reconstructed most of the reference plasmids (recall = 0.82) but approximately a quarter of the predicted plasmid contigs were false positives (precision = 0.76). PlasmidSPAdes merged 83 % of the predictions from genomes with multiple plasmids in a single bin. Recycler searches the assembly graph for sub-graphs corresponding to circular sequences and correctly predicted small plasmids but failed with long plasmids (recall = 0.12, precision = 0.30). cBar, which applies pentamer frequency composition analysis to detect plasmid-derived contigs, showed an overall recall and precision of 0.78 and 0.64. However, cBar only categorizes contigs as plasmid-derived and does not bin the different plasmids correctly within a bacterial isolate. PlasmidFinder, which searches for matches in a replicon database, had the highest precision (1.0) but was restricted by the contents of its database and the contig length obtained from de novo assembly (recall = 0.36).

Surprisingly, PlasmidSPAdes and Recycler detected single isolated components corresponding to putative novel small plasmids (<10 kbp) which were also predicted as plasmids by cBar.

This study shows that it is possible to automatically predict plasmid sequences, but only for small plasmids. The reconstruction of large plasmids (>50 kbp) containing repeated sequences remains challenging and limits the high-throughput analysis of WGS data.

**Author Summary:** Short read sequencing of the DNA of bacteria is often used to understand characteristics such as antibiotic resistance. However the assembly of short read sequencing data with the goal of reconstructing a complete genome is often fragmented and leaves gaps. Therefore independently replicating DNA fragments called plasmids cannot easily be identified from an assembly. Lately a number of programs have been developed to enable the automated prediction of the sequences of plasmids. Here we tested these programs by comparing their outcomes with complete genome sequences. None of the tested programs were able to fully and unambiguously predict distinct plasmid sequences. All programs performed best with the prediction of plasmids smaller than 50 kbp. Larger plasmids were only correctly predicted if they were present as a single contig in the assembly. While predictions by PlasmidSPAdes and cBar contained most of the plasmids, they were merged with or indistinguishable from other plasmids and sometimes chromosome sequences. PlasmidFinder missed most plasmids but all its predictions were correct. Without manual steps or long-read sequencing information, plasmid reconstruction from short read sequencing data remains challenging.

## Introduction

Plasmids are a major driver of variation and adaptation in bacterial populations. The dissemination of multidrug resistance via transfer of plasmids leads to new antibiotic resistant bacteria such as *Escherichia coli* producing extended-spectrum beta-lactamases [1] or vancomycin resistant *Enterococcus faecium* causing nosocomial outbreaks [2]. The prevalence of a plasmid in a bacterial population can increase due to environmental pressures that include the presence of an antibiotic, but may cause a decrease in bacterial fitness in absence of selective pressure [3].

A bacterial cell can hold no, one or multiple plasmids with varying sizes and copy numbers. Traditionally, plasmid sequencing involved the extraction of plasmids using methods to specifically purify plasmid DNA, followed by shot-gun sequencing, which frequently necessitated closing of gaps by PCR or primer-walking [4]. Plasmid DNA purification is exceedingly difficult if it involves plasmids longer than 50 kbp [4, 5]. Alternatively, plasmid sequences can be assembled from whole genome sequencing data (WGS) generated by high-throughput short-read sequencing platforms. However, plasmids often contain repeated sequences shared between the different physical DNA units of the genome, which prohibits complete assembly from short read data. Assembly often results in many fragmented contigs per genome and their origin (plasmid or chromosome) thus remains unclear [6]. Assembly alone is insufficient to determine the origin of a contig and to differentiate contigs belonging to different plasmids. Recently, attempts to reconstruct plasmids from WGS data were automated in a number of programmes.

Currently available plasmid reconstruction programmes either aim to determine whether a previously assembled contig is obtained from a plasmid (PlasmidFinder, cBar), or try to reconstruct whole plasmid sequences from the (mapped) sequencing reads or the assembly graph (Recycler, PlasmidSPAdes, PLACNET) (Table 1).

One of the most widely used tools for plasmid detection and classification is a web tool called PlasmidFinder, developed to detect replicon sequences particularly originating from the family *Enterobacteriaceae* [7]. Two plasmids sharing the same replication mechanism cannot coexist in the long term within the same cell thus replicon sequences are used to classify plasmids into different incompatibility groups [12].

Unsupervised binning using differences in k-mer composition has been widely used in shotgun metagenomic algorithms [13–15]. Composition-based classification methods allow the clustering of contigs into distinct genomes and perform a species-level classification. However, most of these methods are not designed for application to isolated strains and do not report a classification between plasmid or chromosomal contigs. cBar was specifically designed to predict plasmid-derived sequences based on differences in k-mer composition [8]. It relies on differences in pentamer frequencies from 881 complete prokaryotic sequences and gives a binary classification of chromosome-or plasmid-derived contigs.

**Table 1.** Overview of the programmes to reconstruct or predict plasmids from short read sequencing data.

|  | Input | Paired-end information | Coverage | k-mer composition | de Bruijn graph | Similarity to replicons | Similarity to relaxases | Similarity to plasmids | Web-tool | Command-line interface | Included in study |
| --- | --- | --- | --- | --- | --- | --- | --- | --- | --- | --- | --- |
| PlasmidFinder [7] | Contigs |  |  |  |  | ✓ |  |  | ✓ |  | ✓ |
| cBar [8] | Contigs |  |  | ✓ |  |  |  |  |  | ✓ | ✓ |
| Recycler [9] | BAM+assembly graph | ✓ | ✓ |  | ✓ |  |  |  |  | ✓ | ✓ |
| PlasmidSPAdes [10] | Reads | ✓ | ✓ |  | ✓ |  |  |  |  | ✓ | ✓ |
| PLACNET [11] | BAM/SAM+contigs | ✓ | ✓ |  |  | ✓ | ✓ | ✓ |  | ✓ |  |

Plasmid constellation network (PLACNET) reconstructs plasmids from WGS by integrating three lines of evidence: (i) scaffold linking and coverage information from genome assembly, (ii) presence of replication initiator proteins (Rip) and relaxase proteins (Rel), (iii) similarity of the sequences with a custom database containing non-redundant plasmid sequences from NCBI [11]. PLACNET merges all the information into a single network where each component corresponds to a physical DNA unit. Repetitive sequences such as transposases or insertion sequences (IS) with a higher coverage are shared between components. Manual pruning in Cytoscape is necessary to duplicate and split the graph to obtain disjoint components in the final network [16, 17]. Prediction reproducibility rates is thus highly dependent on the expertise of the researcher. As we aimed to test fully automated methods for plasmid reconstruction, we excluded PLACNET from the comparison.

More recently, two algorithms that reconstruct plasmids on basis of the information contained in the *de Bruijn* graph were developed: Recycler [9] and PlasmidSPAdes [10].

Recycler extracts the information from the de Bruijn graph searching for sub-graphs (cycles) corresponding to plasmids. Selection of the cycles is based on the following assumptions: (i) nodes forming a plasmid have a uniform coverage, (ii) a minimal path must be selected between edges because of repetitive sequences, (iii) contigs belonging to the same cycle have concordant read-end paired information and (iv) plasmid cycles exceed a minimum length.

PlasmidSPAdes assumes a highly uniform coverage of the contigs within the chromosome. It calculates the median coverage from the SPAdes assembly graph to estimate the chromosome coverage. By default, only contigs longer than 10 kbp are considered because repeated sequences are mostly present in shorter contigs and long contigs have a lower coverage variance. Contigs are classified as chromosomal edges if their coverage does not exceed a maximum deviation (default 0.3) from the median coverage. PlasmidSPAdes iteratively removes long chromosomal edges to transform the assembly graph into a plasmid graph. Finally, connected components in the plasmid graph are reported as putative plasmids.

Here, we benchmarked currently available programmes to detect and reconstruct plasmid sequences from short read sequencing data, starting either from the reads or from assembled contigs. The aim of this study was to determine whether it is possible to obtain complete plasmid sequences with state-of-the-art tools without manual expert intervention.

**Table 2.** Bacterial genomes included in this study.

| Genome | SRA | Genome accession | Number of plasmids | Range size | Total length |
| --- | --- | --- | --- | --- | --- |
| *Burkholderia cenocepacia* strain DDS 22E-1 | SRR1618480 | GCA_000755725.1 | 0 | - | 0 |
| *Bacillus subtilis* subsp. natto BEST195 | DRR016448 | GCA_000209795.2 | 1 | 5.8 | 5.8 |
| *Enterobacter aerogenes* strain CAV1320 | SRR2965748 | GCA_001021995.1 | 1 | 13.9 | 13.9 |
| *Providencia stuartii* strain ATCC 33672 | SRR1558174 | GCA_000754345.1 | 1 | 48.8 | 48.8 |
| *Corynebacterium callunae* DSM 20147 | SRR892039 | GCA_000420585.1 | 2 | 4.1-85.0 | 89.132 |
| *Enterobacter cloacae* strain CAV1411 | SRR2965820 | GCA_001022075.1 | 2 | 33.6-90.4 | 124.0 |
| *Enterobacter cloacae* strain CAV1669 | SRR2965616 | GCA_001022255.1 | 2 | 33.6-90.4 | 124.0 |
| *Enterobacter cloacae* strain CAV1311 | SRR2965815 | GCA_001022015.1 | 3 | 3.2-90.4 | 127.2 |
| *Enterobacter cloacae* strain CAV1668 | SRR2965612 | GCA_001022055.1 | 2 | 43.4-85.1 | 128.6 |
| *Klebsiella pneumoniae* strain Kpn223 | SRR3465557 | GCA_001663435.1 | 1 | 170.9 | 170.9 |
| *Escherichia coli* JJ1886 | SRR933487 | GCA_000493755.1 | 5 | 1.5-110.0 | 178.3 |
| *Aeromonas veronii* strain AVNIH1 | SRR3465535 | GCA_001634325.1 | 1 | 198.3 | 198.3 |
| *Klebsiella pneumoniae* strain AATZP | SRR3228444 | GCA_001648215.1 | 3 | 38.3-121.0 | 213.4 |
| *Klebsiella pneumoniae* strain CAV1596 | SRR1582868 | GCA_001022235.1 | 4 | 2.9-96.7 | 218.3 |
| *Klebsiella pneumoniae* strain CAV1392 | SRR1582895 | GCA_001022035.1 | 3 | 43.6-130.7 | 224.1 |
| *Escherichia coli* JJ1887 | SRR933489 | GCA_001593565.1 | 5 | 1.5-130.6 | 250.4 |
| *Citrobacter freundii* CFNIH1 | SRR1284629 | GCA_000648515.1 | 1 | 272.2 | 272.2 |
| *Enterobacter asburiae* strain CAV1043 | SRR2965752 | GCA_001022095.1 | 6 | 1.9-96.8 | 278.0 |
| *Klebsiella pneumoniae* strain KPNIH36 | SRR3222156 | GCA_001675125.1 | 3 | 40.44-133.4 | 287.5 |
| *Enterococcus faecium* strain ATCC 700221 | SRR3176159 | GCA_001594345.1 | 3 | 39.1-189.4 | 292.2 |
| *Escherichia coli* strain ECO889 | SRR3465539 | GCA_001663475.1 | 2 | 88.0-212.1 | 300.2 |
| *Klebsiella pneumoniae* subsp. pneumoniae KPNIH24 | SRR1501128 | GCA_000714675.1 | 3 | 58.0-194.8 | 338.4 |
| *Enterobacter cloacae* ECR091 | SRR1576808 | GCA_000750275.1 | 3 | 50.3-176.9 | 338.5 |
| *Serratia marcescens* strain CAV1492 | SRR2965730 | GCA_001022215.1 | 5 | 3.2-199.4 | 351.3 |
| *Citrobacter freundii* strain CAV1741 | SRR2965739 | GCA_001022275.1 | 6 | 1.9-129.1 | 361.1 |
| *Klebsiella pneumoniae* subsp. pneumoniae KPNIH1 | SRR1505904 | GCA_000281535.2 | 3 | 15.0-243.8 | 372.5 |
| *Klebsiella pneumoniae* subsp. pneumoniae KPNIH10 | SRR1427234 | GCA_000281435.2 | 3 | 15.0-243.8 | 372.5 |
| *Klebsiella pneumoniae* strain Kpn555 | SRR3465562 | GCA_001663455.1 | 3 | 26.4-224.4 | 393.7 |
| *Klebsiella oxytoca* strain CAV1099 | SRR2965639 | GCA_001022295.1 | 5 | 5.4-113.9 | 412.8 |
| *Enterobacter cloacae* ECNIH3 | SRR1576778 | GCA_000750225.1 | 4 | 50.3-255.0 | 427.9 |
| *Klebsiella pneumoniae* strain KPNIH39 | SRR3217430 | GCA_001663295.1 | 3 | 36.7-284.8 | 428.1 |
| *Klebsiella oxytoca* strain CAV1335 | SRR2965660 | GCA_001022115.1 | 5 | 5.4-117.6 | 443.5 |
| *Rhodobacter sphaeroides* 2.4.1 | SRR522246 | GCA_000273405.1 | 5 | 52.1-124.3 | 496.7 |
| *Citrobacter freundii* strain CAV1321 | SRR2965690 | GCA_001022155.1 | 9 | 1.9-234.7 | 512.4 |
| *Klebsiella oxytoca* KONIH1 | SRR1501122 | GCA_000714655.1 | 3 | 133.3-205.5 | 532.7 |
| *Klebsiella pneumoniae* strain CAV1344 | SRR1582875 | GCA_001022175.1 | 5 | 3.7-250.3 | 547.9 |
| *Klebsiella pneumoniae* strain CAV1193 | SRR2965672 | GCA_001456135.1 | 5 | 3.7-257.9 | 555.5 |
| *Kluyvera intermedia* strain CAV1151 | SRR2965721 | GCA_001022135.1 | 4 | 43.6-295.6 | 637.7 |
| *Enterobacter cloacae* ECNIH2 | SRR1515967 | GCA_000724505.1 | 3 | 47.2-319.9 | 649.7 |
| *Klebsiella pneumoniae* strain PMK1 | SRR1508819 | GCA_000764615.1 | 4 | 69.9-304.5 | 673.7 |
| *Klebsiella pneumoniae* subsp. pneumoniae KPNIH27 | SRR1427243 | GCA_000695935.1 | 5 | 80.4-338.8 | 890.8 |
| *Klebsiella oxytoca* strain CAV1374 | SRR2965655 | GCA_001022195.1 | 11 | 1.9-332.9 | 969.8 |

## Materials and Methods

### Test data

We selected 42 complete genome sequences with publicly available Illumina Miseq or Hiseq reads (Table 2). All strains were previously sequenced by Pacific Biosystems PacBio RS II and Illumina Miseq or Hiseq with paired-end libraries. Complete genome sequences were downloaded from GenBank and reads from the NCBI Sequence Read Archive (SRA) (Table 2). Low-quality bases at both ends of the reads were trimmed using default paramaters in seqtk (version: 1.0-r31, https://github.com/lh3/seqtk.git).

### Plasmid prediction

We predicted plasmids from short reads with four different programs: PlasmidFinder, cBar, Recycler and PlasmidSPAdes. *De novo* assembly was performed using SPAdes 3.8.2 on a high performance computer running CentOS7. For each sample, the assembly graph and resulting contigs corresponding to the maximum *k-mer* used by SPAdes 3.8.2 were selected [18]. Contigs with a size less than 500 bp were filtered out.

- *PlasmidFinder*. To replicate results that would be obtained through the use of the PlasmidFinder web interface, we downloaded the PlasmidFinder database containing 121 replicon sequences (updated on 16 March 2016) from the Center for Genomic Epidemiology (https://cge.cbs.dtu.dk//services/data.php). We then performed nucleotide BLAST (NCBI-BLAST version 2.2.28+) searches against this database [19]. Contigs were identified as plasmids if they had a minimum identity of 80% and covered at least 60% of the replicon sequence, consistent with the parameters used to identify plasmids in bacterial whole-genome data by the authors of PlasmidFinder [7]. Contigs in which a replicon sequence was identified were considered as PlasmidFinder prediction.
- *cBar*. We downloaded cBar version 1.2 from http://csbl.bmb.uga.edu/ffzhou/cBar/cBar.1.2.tar.gz and used it to categorize contigs derived by SPAdes 3.8.2. Contigs categorized as plasmid-derived were considered as cBar prediction.
- *Recycler*. We downloaded Recycler (single version, date: 07-03-2016) from https://github.com/Shamir-Lab/Recycler. The BAM file required as input by Recycler was created by alignment of the trimmed reads against the resulting contigs using Bwa 0.7.12 [20] and samtools 1.3.1 [21]. Cycles reported in the assembly graph were considered as Recycler prediction.
- *PlasmidSPAdes*. We run PlasmidSPAdes (packaged in SPAdes 3.8.2) with standard parameters. The components reported in contigs.fasta were considered as PlasmidSPAdes prediction.

## Measures for the evaluation

We evaluated the performance of each programme regarding accuracy and completeness. Quast (version 4.1) [22] was used to map plasmid predictions against i) each reference plasmid separately or ii) the reference genome (containing chromosomes and plasmids) using Nucmer alignments. Total contig length was used to estimate the following terms:

- **Plasmid fraction**. Fraction of the prediction that matched the reference plasmids (true positive prediction). Due to the presence of repeated sequences, contigs can map to both the reference plasmids and the chromosome. These contigs were scored as true positives and only included within the plasmid fraction.
- **Chromosome fraction**. Fraction of the prediction that matched the reference chromosome (false positive prediction). This fraction can include non-plasmid mobile genetic elements from the chromosome such as phages or transposable elements.
- **Fraction of novel sequences**. Fraction of the prediction not mapping to either the reference plasmid or the chromosome, thus corresponding to contigs absent from the reference assembly.

Novel contigs were further analyzed and annotated using Prokka (version 1.12-beta) [23]. To identify potential novel plasmids we compared these sequences to the non-redundant nucleotide database of the NCBI using BLAST. The best blast hit was extracted selecting minimum e-value and highest bit-score as previously described [10]. Furthermore, the completeness of the potential novel mobile genetic elements was corroborated by generating a dot-plot aligning the sequence to itself [24]. The presence of the same repeated sequence at the ends of the contig suggested a potential circularization signature. This analysis was summarized in Table 3.

The programs were further evaluated using the following metrics.

- **Recall** was defined as the percentage of the reference plasmid(s) covered by the prediction. On the individual plasmid level, a recall of 100% indicates that the full sequence of the reference plasmid was present among the predicted plasmids. On the whole genome level, a recall of 100% indicates all reference plasmids were fully present among the predicted plasmids. However, recall does not take prediction of plasmid synteny or plasmid boundaries into account.

Recall value was estimated using the genome fraction reported in Quast.

- **Precision**. We defined precision as:

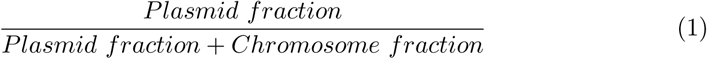

The *fraction of novel sequences* was ignored when calculating precision.

The negative control, the plasmid-less *B. cenocepacia* strain 22E-1, was excluded from recall and precision calculations. Icarus [25] (packaged in Quast 4.1) was used to visualize the alignments between the reference genomes and the predicted sequences. Scaffold linkage of specific contigs in the PlasmidSPAdes assembly graph of a selection of genomes was visualized with Bandage (version 0.7.1) [26].

The workflow (S1 Fig) was written in bash and python (version 2.7) and subsequent analysis in R (version 0.99.982). Scripts and a detailed explanation of the analysis are available as a git repository at (git@gitlab.com:sirarredondo/Plasmid_Assembly.git).

## Results

### Reference genomes

The test data included sequences of complete bacterial genomes from twelve different genera: *Aeromonas, Bacillus, Burkholderia, Citrobacter, Corynebacterium, Enterobacter, Escherichia, Klebsiella, Kluyvera, Providencia, Rhodobacter* and *Serratia*. In total, the test data contained 148 plasmid sequences ranging from 1.55 to 338.85 kbp (Figure 1) and 45 chromosomal sequences ranging from 0.93 to 6.26 Mbp.

The most complex composition of plasmids was present in *K. oxytoca* strain CAV1374 with a single chromosome and eleven plasmids ranging from 1.91 to 332.95 kbp (Figure 1). In contrast, *B. subtilis* subsp. natto BEST195 contained a single plasmid with a length of 5.84 kbp (Figure 1).

*B. cenocepacia* DDS 22E-1 was included as a negative control as this strain does not carry plasmids (Figure 1), but contains three chromosomes with a length of 1.17, 3.21 and 3.67 Mbp. In addition, *R. sphaeroides* 2.4.1 contained two chromosomes with a length of 3.19 and 0.94 Mbp along with 5 plasmid ranging from 52.1 to 124.3 kbp.

**Fig 1.**
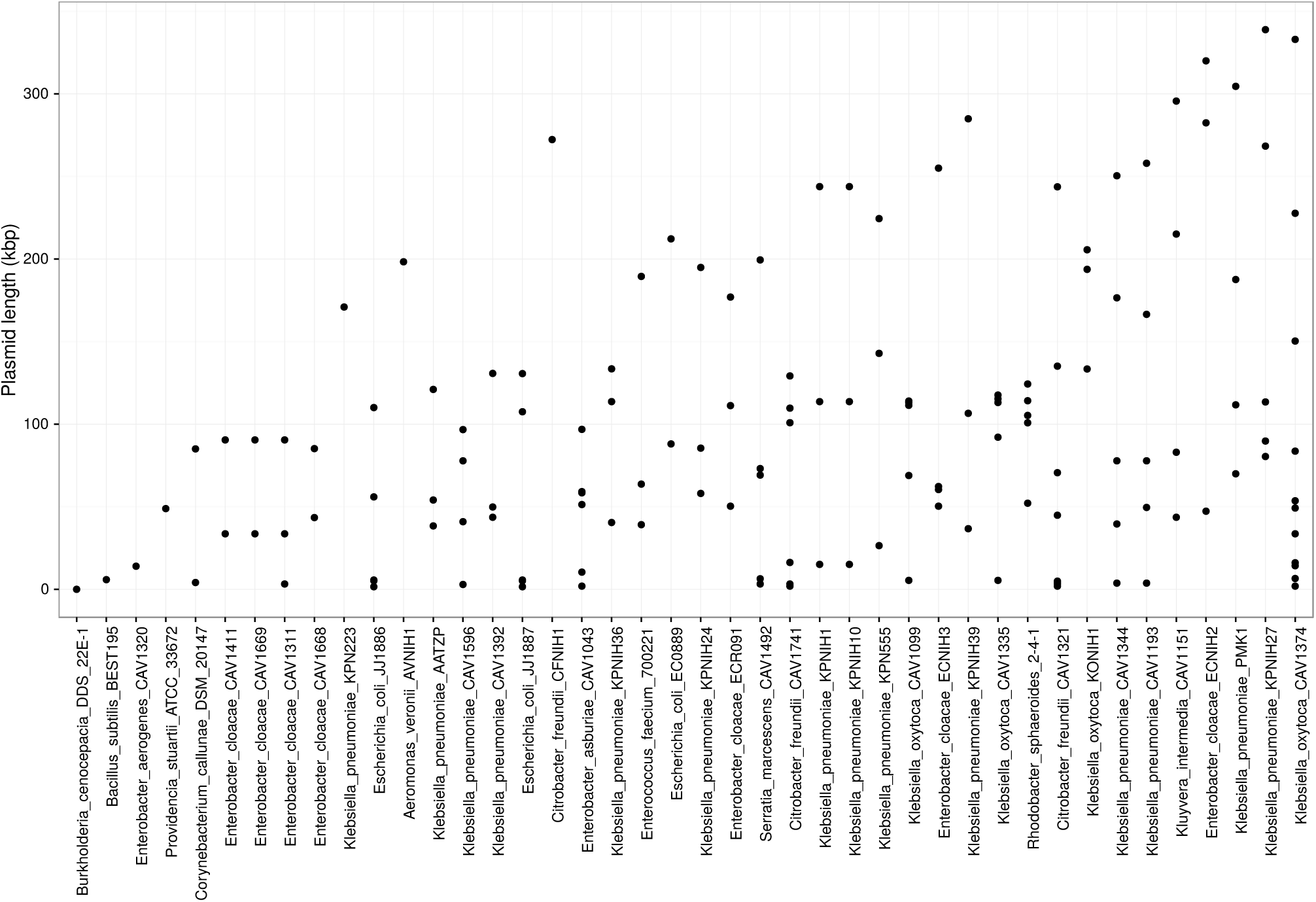
Overview of reference plasmids. The size of the reference plasmids is shown for each bacterial isolate. Strains were sorted based on their total plasmid length. *B. cenocepacia* DDS 22E-1 was considered as a negative control because no reference plasmids are present. *K. oxytoca* strain CAV1374 was the most complex isolate with eleven plasmids ranging from 1.9 to 332.9 kbp

### Reconstruction per plasmid

The performance of the programs was first evaluated on a single plasmid level. We defined a minimum recall value of 0.9 to classify a plasmid as correctly predicted. Out of 148 reference plasmids included in this study, 133 (89.9 %) were reconstructed by either PlasmidFinder, cBar, Recycler or PlasmidSPAdes (Figures 2 and 3). PlasmidSPAdes recovered 125 plasmids, cBar 84 plasmids, Recycler 21 plasmid and PlasmidFinder 13 plasmids at a recall of 0.9 or more (Figure 3). While the recall value of reference plasmids by the predictions declined with plasmid size for Recycler, cBar and PlasmidFinder predictions, the predictions of PlasmidSPAdes were not affected by plasmid length. Recall values obtained for each reference plasmid are available at S1 Table

Five genomes (*E. coli* JJ1886; *R. sphaeroides* 2.4.1, *C. freundii* CFNIH1, *B. cenocepacia* strain DDS 22E-1 and *C. callunae* DSM 20147) were previously used to validate Recycler and/or PlasmidSPAdes [9, 10]. Recall values obtained for each of the reference plasmids in this study were concordant with previous findings (S1 Table and S1 Appendix).

Of all 148 plasmids, five plasmids were consistently fully predicted by all of the programs (Figure 3). These included two large plasmids of 109.6 and 111.6 kbp belonging to the bacterial isolates *K. pneumoniae* strain CAV1392 and *K. pneumoniae* strain PMK1. Visualization of the contigs alignments showed that these plasmids were fully assembled in a single SPAdes contig which did not have any similarity to other reference plasmids or chromosome of the same bacterial genome. In contrast, 15 plasmids consistently had a recall value less than 0.9 in all predictions. Four of these 15 plasmids were not fully covered by SPAdes contigs, therefore precluding complete assembly of the plasmids.

**Fig 2.**
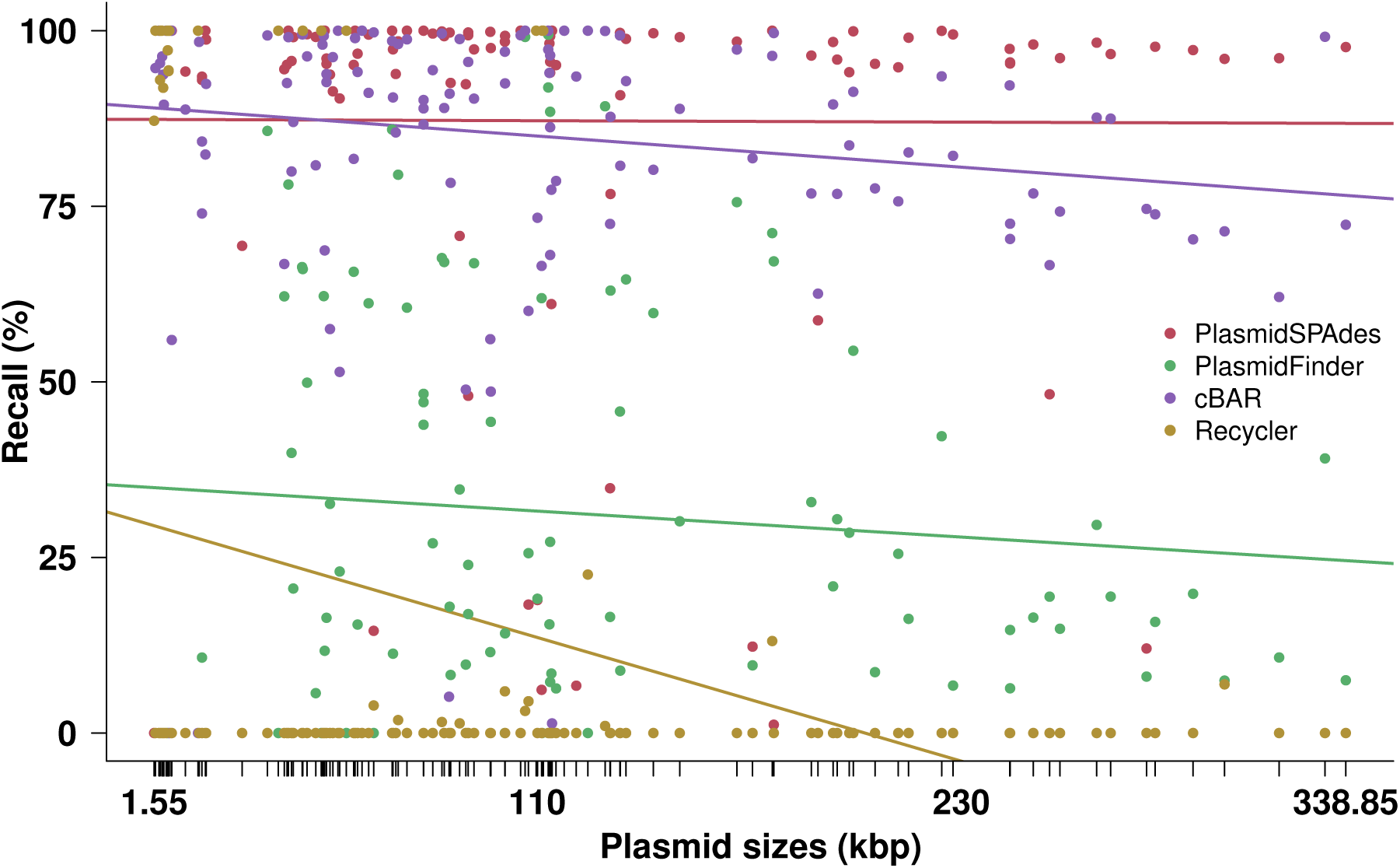
Recall of reference plasmids by predicted plasmid sequences from PlasmidSPAdes, PlasmidFinder, cBar and Recycler. Recall was calculated by aligning the reference plasmid sequences against the plasmid predictions of each genome and disregarded plasmid binning (if any). Lines indicate linear least squares regression fits to data points. Tick marks on the x-axis represent plasmid sizes.

The definition of recall per plasmid operated here does not take into account if plasmid boundaries were called correctly. Both programmes with a high average recall (PlasmidSPAdes and cBar, 0.87 and 0.86, respectively) did not, or incompletely, report plasmid boundaries. cBar performs a binary classification predicting contigs as either “plasmid” or “chromosome” but did not sort the sequences into different plasmids from the same bacterial isolate. PlasmidSPAdes merged plasmids in 83 % of all the genomes with several reference plasmids, and plasmid boundaries were not readily retrievable.

### Reconstruction per genome

Next, performance was evaluated on the genome level, thus comparing the entirety of all predicted plasmid sequences of each genome against all (1-11) plasmids of each genome per programme. We assigned each as plasmid predicted contig to one of the following three categories: (i) plasmid fraction, (ii) chromosome fraction, (iii) novel sequences fraction. Subsequently, precision and recall values were calculated.

**Fig 3.**
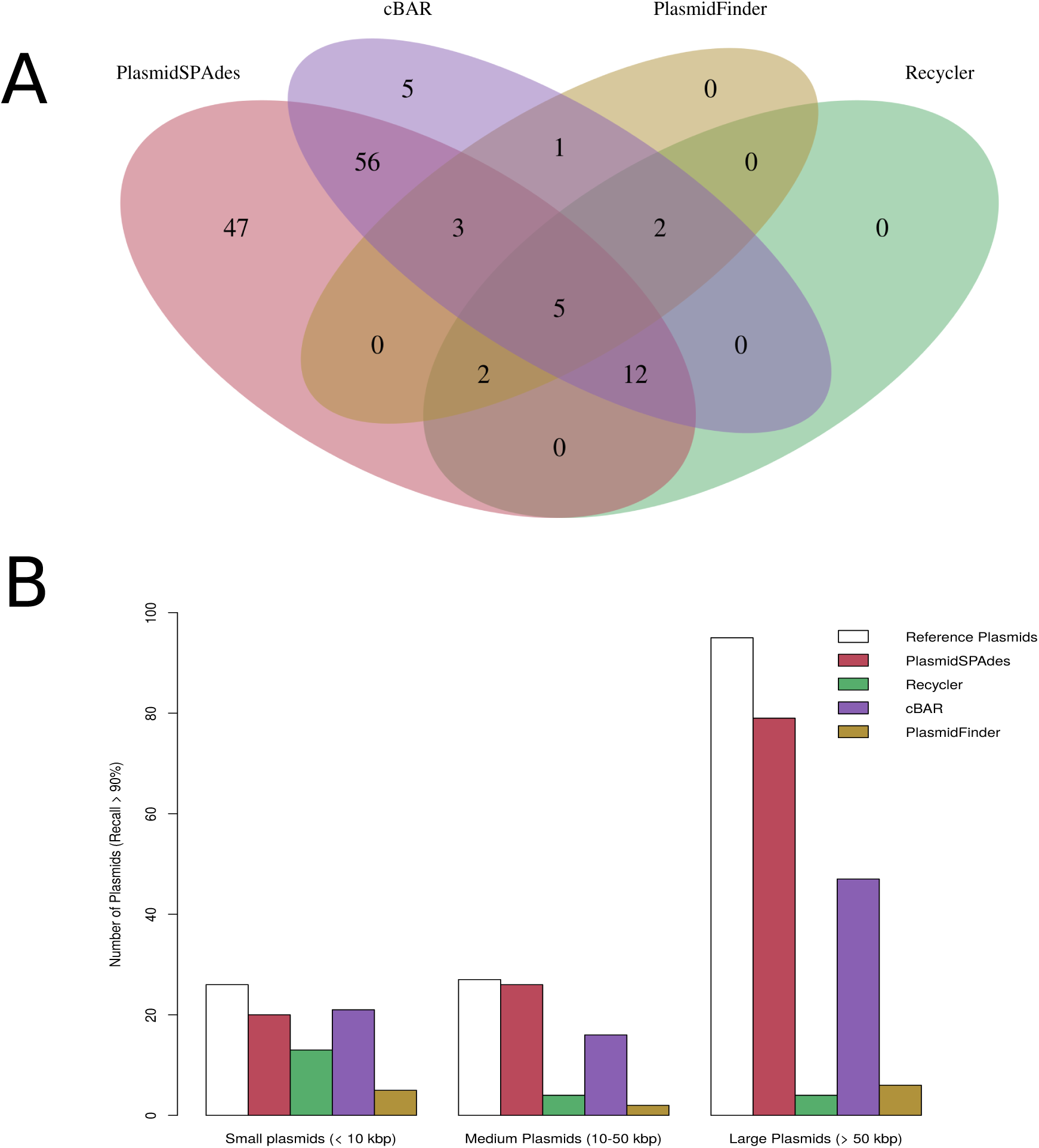
Performance of the programs on single plasmid level. A minimum recall value of 90 % in the program prediction was selected to consider a plasmid as correctly reconstructed. **A**. Venn diagram showing the overlap in prediction between PlasmidSPAdes (red), cBar (purple), PlasmidFinder (orange) and Recycler (green). The intersection of the ellipses showed five plasmids present in all the predictions. **B**. Reference plasmids were classified into small (less than 10 kbp), medium (from 10 to 50 kbp) and large plasmids (greater than 50 kbp) depending on their size. The number of reference plasmids correctly predicted by the programs is represented in the three categories.

**Fig 4.**
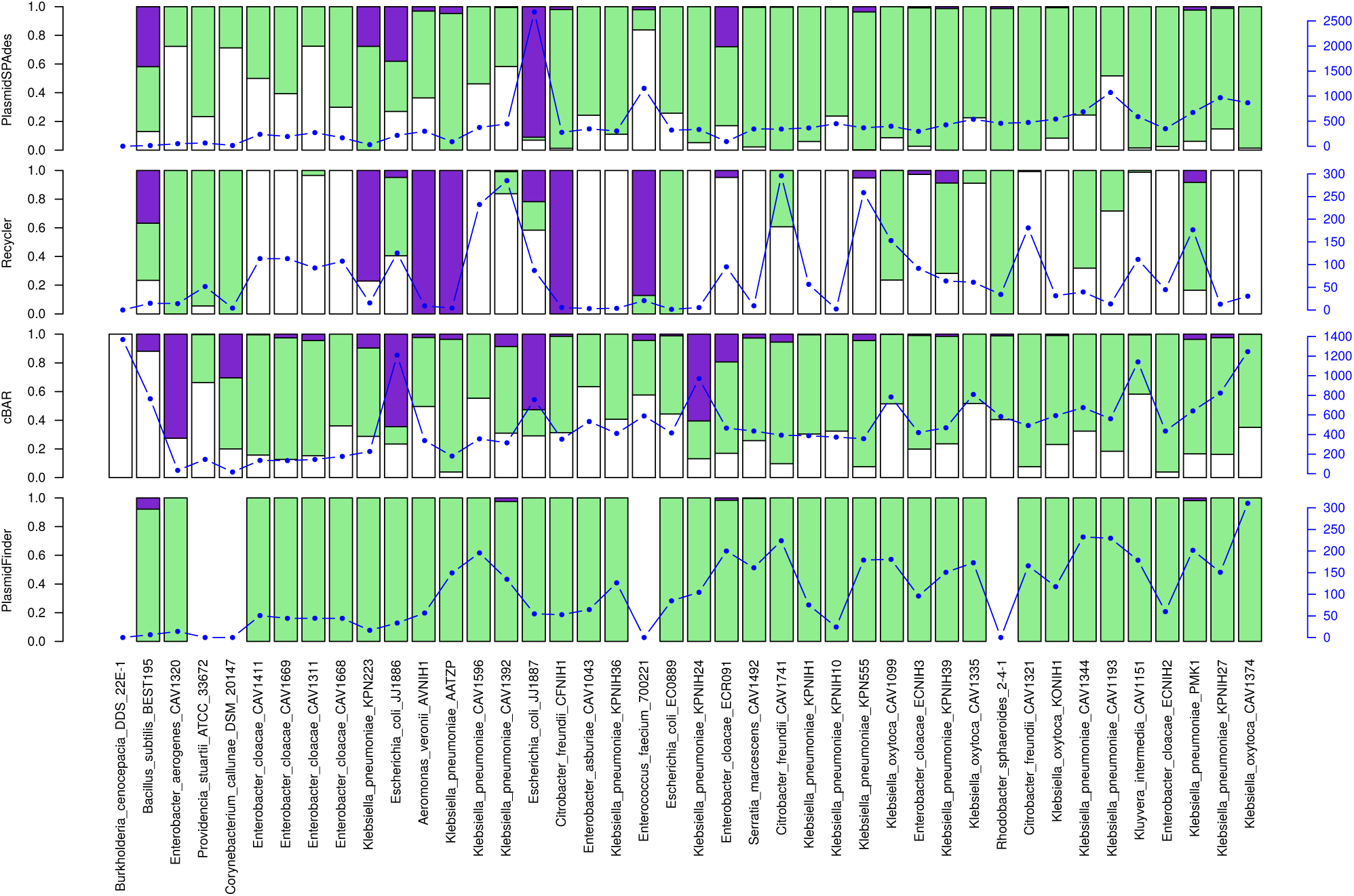
Performance of the programmes on genome level. The prediction of each program was mapped against the reference genomes of each bacterial isolate. Contigs mapping to the reference plasmids were depicted as plasmid fraction (green bar), to the reference chromosome as chromosome fraction (white bar) or to neither as novel sequences fraction (purple bar). On the right y-axis the total length (in kbp) of reconstructed plasmid contigs is indicated. cBar was the only program predicting contigs as plasmids in the genome that was used as negative control (*B. cenocepacia* DDS 22E-1).

### PlasmidSPAdes

PlasmidSPAdes obtained an average plasmid fraction of 0.72 and an average chromosome fraction of 0.22 (Figure 4). Surprisingly, a fraction of 0.06 corresponding to contigs not mapping to the reference genomes was detected. This resulted in an overall precision of 0.76. The majority of plasmids were present in the prediction (overall recall = 0.82).

The completeness of the prediction was high even in the bacterial isolates with an elevated number of reference plasmids. However, PlasmidSPAdes merged plasmid contigs into a single bin when they shared repeated sequences. For example, *K. oxytoca* strain CAV1374 contained eleven reference plasmids and most of these were predicted (recall value = 0.97, Figure 5). However, PlasmidSPAdes merged all the predicted contigs into a single bin with a size of 870.8 kbp. In total, 14 contigs with a size ranging from 0.6 to 4.7 kbp matched to two or more reference plasmid sequences from the *K. oxytoca* strain CAV1374, among which a transposase with a length of 3.6 kbp present in six reference plasmids.

**Fig 5.**
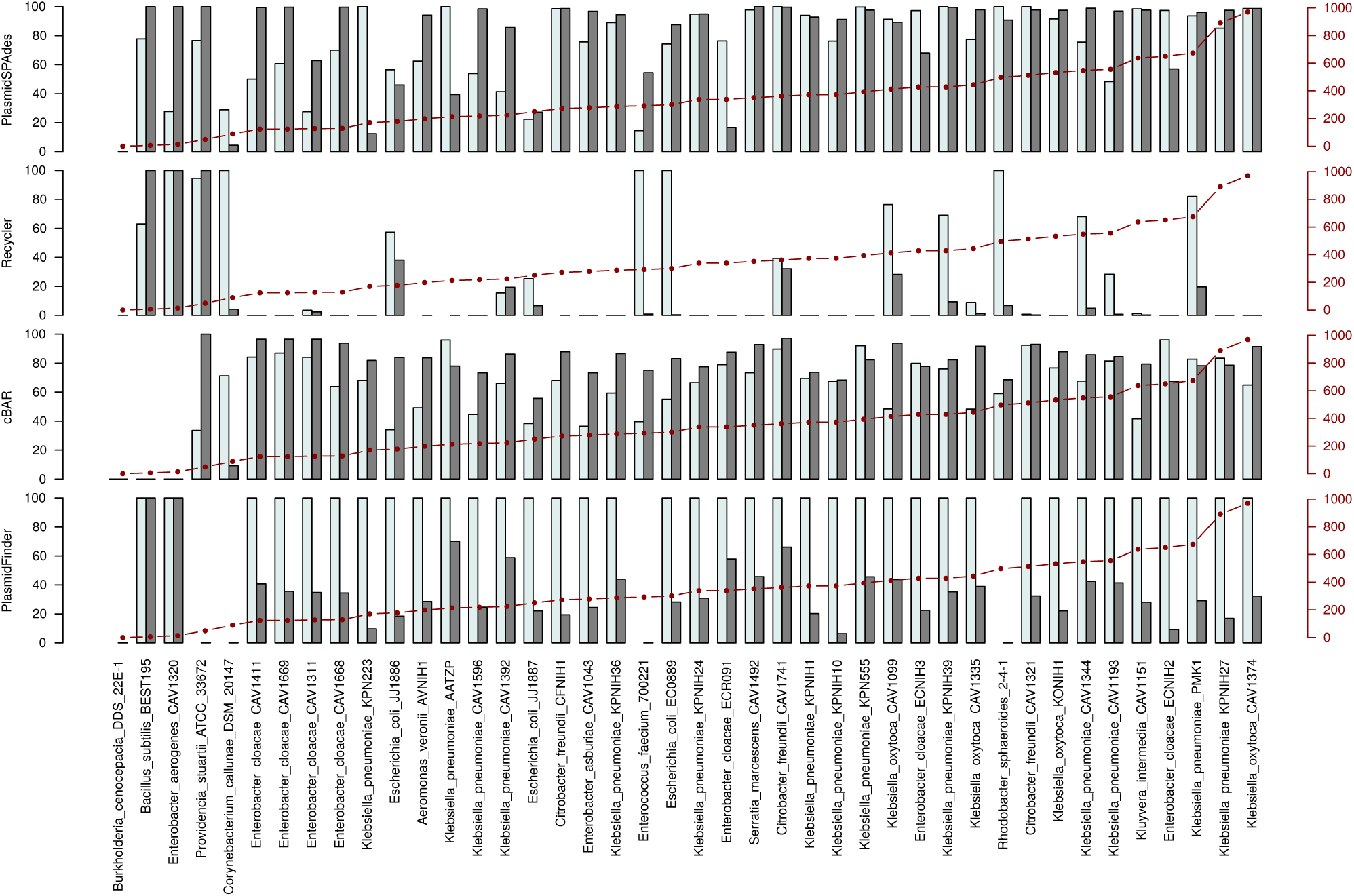
Precision and recall values for each bacterial genome. Precision and recall values are represented (in percentage) with white and gray bars respectively. A precision of 100% indicates the absence of contigs mapping to the reference chromosome in the prediction. Recall of 100% indicates the full sequences of all the reference plasmids was present in the prediction. On the right y-axis is indicated (in kbp) the total plasmid length of a particular bacterial genome.

PlasmidSPAdes predicted a total contig length of 18.18 Mbp of which 3.78 Mbp were mapping to the reference chromosome. These chromosomal contigs were analyzed and annotated by Prokka to search for phage-related genes. We applied the same keywords as reported in Phaster [27, 28] to assess the presence of a phage sequence. A total of 1.36 Mbp showed evidence for presence of phage-related genes. These sequences were possibly predicted as plasmids because their coverage differed from the host genome.

We found contigs not mapping to the reference genomes in 20 cases (Figure 4). With the exception of *E. coli* JJ1886 and *E. coli* JJ1887, most of the contigs present in the fraction of novel sequences were detected as isolated components by PlasmidSPAdes. Copy number of those components was inferred from their k-mer coverage ratio. Isolated components including a single contig are highlighted in Table 3. Contig size, best blast hit, inferred copy number, gene annotation and circularity are reported.

**Table 3.** Novel sequences not present in the reference genome predicted by PlasmidSPAdes and Recycler.

|  | pSPAdes | Recycler | cBar | Copy number | Blast hit | Annotation | Circularity |
| --- | --- | --- | --- | --- | --- | --- | --- |
| *B. subtilis* BEST195 | 5513 | 5386 | Plasmid | 10.3 | Plasmid (CP003995) | - | ✓ |
| *K. pneumoniae* KPN223 | 4294 | 4167 | Plasmid | 0.9 | Plasmid (EU932690) | - | ✓ |
|  | 4141 | 4014 | Plasmid | 1.5 | Non significant | Mob. protein MobA | ✓ |
|  | - | 3478 | Plasmid | 1.3 | Plasmid(NZ CP012489) | - | ✓ |
| *E. coli* JJ1886 | 11105 | - | Chromosome | 0.2 | Chromosome (CP013218) | - | X |
|  | - | 2361 | Plasmid | 1.0 | Plasmid (CP014694) | - | ✓ |
|  | - | 2216 | Plasmid | 0.8 | Plasmid (Y16944) | - | ✓ |
|  | 1689 | 1634 | Plasmid | 0.2 | Plasmid (JQ312422) | - | ✓ |
| *A. veronii* AVNIH1 | 7241 | 7114 | Plasmid | 6.3 | Plasmid (KT781681) | Antitoxin RelB | ✓ |
|  | 1863 | 1736 | Chromosome | 15.7 | Plasmid (LN853312) | - | ✓ |
| *K. pneumoniae* AATZP | 4294 | 4167 | Plasmid | 2.4 | Plasmid (CP003995) | - | ✓ |
| *K. pneumoniae* CAV1392 | 2572 | 2495 | Plasmid | 0.1 | Plasmid (NC 015515) | - | ✓ |
| *C. freundii* CFNIH1 | 5487 | 5410 | Plasmid | 14.1 | Plasmid (NZ_CP011613) | Relaxase MbeA | ✓ |
| *E. faecium* ATCC 700221 | 12589 | 12462 | Plasmid | 2.7 | Plasmid (AB158402) | - | ✓ |
|  | 5513 | 5386 | Plasmid | 26.6 | Phage (CP004084) | - | ✓ |
| *K. pneumoniae* KPN555 | 4175 | 4048 | Plasmid | 0.4 | Plasmid (JX238446) | Relaxase MbeA | ✓ |
|  | 3605 | 3478 | Plasmid | 0.9 | Plasmid (CP000652) | Antitoxin MazE | ✓ |
|  | 3001 | 2874 | Plasmid | 1.7 | Plasmid (HG796369) | Plasmid recombination enzyme | ✓ |
|  | 2925 | 2798 | Plasmid | 2.0 | Plasmid (HG796369) | - | ✓ |
| *K. pneumoniae* PMK1 | 5695 | 5640 | Plasmid | 26.0 | Plasmid (LN854314) | Antitoxin IgA-2, Mob. protein MbeC | ✓ |
|  | 5441 | 5386 | Plasmid | 2.0 | Scaffold (LL266921) | - | ✓ |
|  | 3825 | 3770 | Plasmid | 35.0 | Plasmid (NC 019077) | - | ✓ |
| *E. cloacae* ECR091 | 4744 | 4667 | Plasmid | 11.8 | Plasmid (CP004060) | Mob. protein MbeC | ✓ |
|  | 2572 | - | Plasmid | 22.0 | Plasmid (AF014880) | - | ✓ |
| *E. cloacae* ECNIH3 | 2572 | 2495 | Plasmid | 30.9 | Plasmid (AF014880) | - | ✓ |
| *K. oxytoca* KONIH1 | 3713 | - | Chromosome | 40.7 | Plasmid (CP011586) | - | ✓ |
| *K. pneumoniae* KPNIH39 | 5550 | 5521 | Plasmid | 9.1 | Plasmid (NC 019346) | - | ✓ |

The fraction of novel sequences reported in *E. coli* JJ1886 and *E. coli* JJ1887 (Figure 4) suggested that contaminants may interfere with plasmid reconstruction by PlasmidSPAdes. Sequences not present in the reference genome had high similarity with chromosome and plasmids of *Staphylococcus aureus* (S1 Appendix). The chromosome and plasmids of *S. aureus* were not filtered out by PlasmidSPAdes because their coverage differed from the host chromosome. Further discussion on the identification of potential novel small cryptic plasmids is available at S2 Appendix.

### Recycler

Recycler obtained an average plasmid fraction of 0.24, an average chromosome fraction of 0.62 and an average fraction of novel sequences of 0.14 (Figure 4). This resulted in an overall precision of 0.30 indicating a high number of sequences originating from the chromosome.

Recycler obtained a low overall recall of 0.12 (Figure 5). This value can partly be explained by the fact that the algorithm only reports unique circular sequences. Therefore plasmids sharing highly similar sequences with each other were not present in the prediction.

The recall value obtained by Recycler was 1.0 in samples with small or medium size plasmids (e.g. *B. subtilis* BEST195 or *E. aerogenes* CAV1320). Furthermore, large plasmids not sharing any repeated sequence with other replicons were also correctly predicted by Recycler. This includes two plasmids of 100.8 and 111.3 kbp from *C. freundii* CAV1741 and *K. oxytoca* CAV1099.

Recycler predicted a total contig length of 3.06 Mbp, of which 2.25 Mbp were mapping to the reference chromosomes. These chromosomal contigs were annotated to detect phage sequences. A total of 1.74 Mbp showed evidence for the presence of phage-related genes. Recycler was designed to extract circular sequences from the assembly graph. Therefore, Recycler predictions also contained non-plasmid mobile genetic elements with a potential circularization signature. The same phage sequence of 41.9 kbp was predicted in *E. cloacae* strain CAV1311, *E.cloacae* strain CAV1411, *E.cloacae* strain CAV1668 and *E.cloacae* strain CAV1669. In most of these isolates, Recycler precision was 0.0 because no reference plasmid sequence was recovered (Figure 5).

Recycler more robustly detected plasmid sequences in contaminated samples than PlasmidSPAdes. In contrast to PlasmidSPAdes, the fraction of novel sequences was not higher in *E. coli* JJ1886 and *E. coli* JJ1887 compared to the rest of genomes (Figure 4).

Most of the novel contigs reconstructed by Recycler were also predicted by PlasmidSPAdes as isolated components. However, in all cases, Recycler trimmed one of the adjoining regions from the circular sequence, reporting the correct plasmid size (Table 3). Common features of these novel contigs are a length less than 10 kbp and an intermediate copy number (S2 Appendix).

### cBar

cBar predicted every contig as either plasmid-derived or chromosome-derived. In order to maintain comparability for recall and precision calculation, we only considered contigs predicted as plasmids. The total size of sequences predicted as plasmids was 21.66 Mbp.

cBar obtained an average plasmid fraction of 0.58, an average chromosome fraction of 0.33 and an average fraction of novel sequences of 0.09. This resulted in an overall precision and recall of 0.64 and 0.78 respectively.

A substantial amount of contigs corresponding to reference plasmids was recovered. The completeness of the results was high despite of the complexity of the sample. For instance, cBar obtained a recall value of 0.93 in *C.freundii* CAV1321 which contained nine reference plasmids (Figure 1). However, the precision varied largely across genomes, as reflected in *P. stuartii* ATCC 33762 which contains a single reference plasmid of 48.87 kbp (Figure 1). This plasmid was correctly detected by cBar resulting in a recall value of 1.0. Nevertheless, it wrongly predicted 19 contigs (>500 bp) as plasmids which mapped to the chromosome, resulting in a precision of 0.34 (Figure 5).

In *B. subtilis* subsp. natto BEST195 and *E. aerogenes* CAV1320, which carry single plasmids, precision and recall value were 0.0 (Figure 5). Both these plasmids were assembled into single contigs but the algorithm erroneously predicted these as chromosome-derived.

Notably, in the negative control *B. cenocepacia* DDS 22E-1, cBar predicted a total size of 1369 kbp wrongly as plasmid-derived contigs 4.

Using cBar, the detection of novel plasmids is more difficult compared to Recycler or PlasmidSPAdes because graph component information is not available. However, only with the exception of two putative small cryptic plasmids in *A. veronii* AVNIH1 and *K. oxytoca* KONIH1, all components highlighted in Table 3 were also classified as plasmids by cBar.

### PlasmidFinder

The total size of contigs with a replicon sequence detected by PlasmidFinder was 4.39 Mbp. PlasmidFinder obtained an average plasmid fraction of 0.99 and an average fraction of novel sequences of 0.01.

PlasmidFinder was able to detect at least one plasmid replicon sequence in 37 bacterial strains, but failed to detect any replicon sequence in *R. sphaeroides* 2-4-1 and in the gram-positive bacteria *C. callunae* DSM 20147, *E. faecium* ATCC 700221 and *P. stuartii* ATCC 33672.

The database of PlasmidFinder was designed to detect replicon sequences from the family *Enterobacteriaceae*. For this reason, we excluded all gram-positive genomes from precision and recall calculations. Surprisingly in *B. subtilis* BEST195, one of the four gram-positive strains, a recall of 1.0 was obtained. Nucleotide blast showed that the single reference plasmid present in *B. subtilis* BEST195 had an identity of 88% and covered 82% of a replicon sequence (NC_015392) from *Salmonella enterica* strain 853 that was indexed in PlasmidFinder database.

The overall precision of PlasmidFinder was 1.0, indicating that no false positive sequences were predicted as plasmids. However, the overall recall of 0.36 was due to the low completeness of the results as shown in Figure 5. The recall of PlasmidFinder was directly linked to the size of the contigs where the replicon sequence was detected. For example, in *B. subtilis* BEST195 and in *E. aerogenes* CAV1320 we obtained a recall value of 1.0 because the strains carried single plasmids with a size of 5.8 and 14.0 kbp respectively. These plasmids were completely assembled into a single SPAdes contig which contained a replicon sequence.

## Discussion

We compared four different programmes to reconstruct or predict plasmid sequences from WGS data. The large majority of the sequences of the plasmids (89.9 %) could be reconstructed by one of the programmes when compared to the reference plasmids. However, in many cases, the reconstructions were fragmented (all programmes), contaminated by chromosome sequences (cBar, Recycler, PlasmidSPAdes), boundaries of the plasmids were unclear (cBar, PlasmidSPAdes) and plasmids incomplete (all programmes). In absence of reference plasmid sequences, disentangling or binning the reconstructions into separate plasmids is a challenging step that still has to be solved.

The overall recall obtained by PlasmidSPAdes (0.82) showed that most of the reference plasmids were fully or partially present in the plasmid prediction. The major drawback in using PlasmidSPAdes was the lack of boundaries when reference plasmids shared a high number of similar sequences. This limitation can be overcome by applying the same methodology as already established in PLACNET [11]. By visualizing the plasmid graph and connecting contigs with a similar coverage and scaffolding linkage, plasmid sub-graphs can be separated manually, if the different plasmids sufficiently differ in their copy number [10] (S1 Appendix). Repeated sequences such as transposases merging different components in the graph can be spotted by their high number of scaffolding links and coverage. To disentangle the graphs it is necessary to assign them to each of the sub-components separately. However, whether manual interventions are successful, is highly dependent on the expertise of the individual analyzing the data, can be difficult to reproduce independently and limits the high-throughput analysis of WGS data.

Recycler applies an innovative and general approach to plasmid reconstruction and successfully extracted complete plasmid sequences if they had circular features present in the assembly graph. Most large plasmids, however, tend to be assembled into several contigs due to the presence of repeated sequences with high coverage. Recycler failed to extract these types of plasmids and in many cases only extracted non-plasmid mobile elements.

cBar was originally designed to categorize chromosome and plasmids in metagenomic sequences by comparing pentamer frequencies of a plasmid database. The accuracy of this approach is known to be lower for long plasmids because the nucleotide composition of these plasmids is similar to the host chromosome [29]. However, the overall recall of cBar is high (0.78) and it might be well-suited to confirm if a sequence is predicted to be plasmid-derived by another method.

The results of PlasmidFinder showed an outstanding 1.0 true positive rate indicating a high reliability of the prediction. Being initially designed for *Enterobacteriaceae*, it was not able to detect any plasmid replication initiator protein in four bacterial strains including three gram-positive genomes. If applied to PlasmidSPAdes predictions, the detection of different incompatibility groups by PlasmidFinder could indicate the presence of two or more plasmids merged together into a single component.

To our surprise, PlasmidSPAdes and Recycler predicted a large number of contigs (fraction of novel sequences: 0.06 and 0.14, respectively) that were not present in the complete reference genomes and which were also predicted as plasmids by cBar. These sequences could originate from sequence reads that were filtered in the reference assembly because they were considered to be contaminant sequences, but could also represent small replicons. As described elsewhere, hierarchical genome assembly process (HGAP) of PacBio reads can lead to missing small plasmids in the main assembly when using a seed read length cut-off greater than actual plasmid size [30, 31]. To include small plasmids in a genome assembly we suggest to perform a subsequent *de novo* assembly using short-reads not mapping to the PacBio assembly or to perform a HGAP iteratively reducing the seed read length when assembling whole genomes.

Small cryptic plasmids are mostly composed of genes involved in plasmid replication and were previously described in ESBL-producing *E. coli* [32]. We analyzed a total of 27 putative small cryptic plasmids extracted either by Recycler or PlasmidSPAdes corresponding to isolated components with a single contig. However experimental validation is required to confirm these plasmids as stable residents.

To obtain the full sequences of plasmids, long read sequencing data can be a solution [33]. Nonetheless, the relatively high error rate of long read sequencing by Pacific Biosystems PacBio RS II or Oxford Nanopore Technologies Ltd makes desirable the combination of long and short-read sequencing technologies for accurate plasmid sequencing. Moreover, we showed the importance of checking the presence of small replicons not present in the reference assembly. This may be crucial to identify the entirety of the plasmids repertoire and, with that, obtain complete genome sequences.

In this study, plasmid reference sequences were present for comparison, something which is lacking in WGS projects for which these tools have been developed. The presence of repeated sequences shared in different physical DNA units, indiscriminate pentamer frequencies and similar coverage ratios make the *de novo* reconstruction of plasmids from WGS challenging, even with the help of the reconstruction programmes tested here.

## Supporting Information

S1 Fig. Analysis workflow.

S1 Appendix. Genomes considered as positive controls.

S2 Appendix. Potential novel small cryptic plasmids.

S1 Table. Recall values for each reference plasmid.

## Acknowledgments

We would like to thank Dimitry Antipov and Roye Rozov for their valuable and insightful comments.

